# Lipid bicelles in the study of biomembrane characteristics

**DOI:** 10.1101/2022.11.23.517649

**Authors:** Matthias Pöhnl, Christoph Kluge, Rainer A. Böckmann

## Abstract

Simulations of lipid membranes typically make use of periodic boundary conditions to mimic macroscopically sized membranes and allow for comparison to experiments performed e.g. on planar lipid membranes or on unilamellar lipid vesicles. However, the lateral periodicity partly suppresses membrane fluctuations or membrane remodeling, processes that are of particular importance in the study of asymmetric membranes – i.e. membranes with integral or associated proteins and/or asymmetric lipid compositions.

Here, we devised a simple albeit powerful lipid bicelle model system that (i) displays similar structural, dynamical and mechanical properties compared to infinite periodic lipid membrane systems, and allows (ii) for the study of asymmetric lipid bilayer systems, and (iii) the unperturbed formation of local spontaneous curvature induced by lipids or proteins in coarse-grained and all-atom molecular dynamics simulations. In addition, the system is characterized by largely unbiased thermal fluctuations as opposed to standard bilayer systems. Application of the bicelle system for an asymmetric lipid composition resembling the plasma membrane reveals that the cholesterol density for a tension-free plasma membrane with a vanishing spontaneous curvature is larger by 28% within the extracellular leaflet compared to the cytosolic leaflet.

**Graphical TOC Entry:** 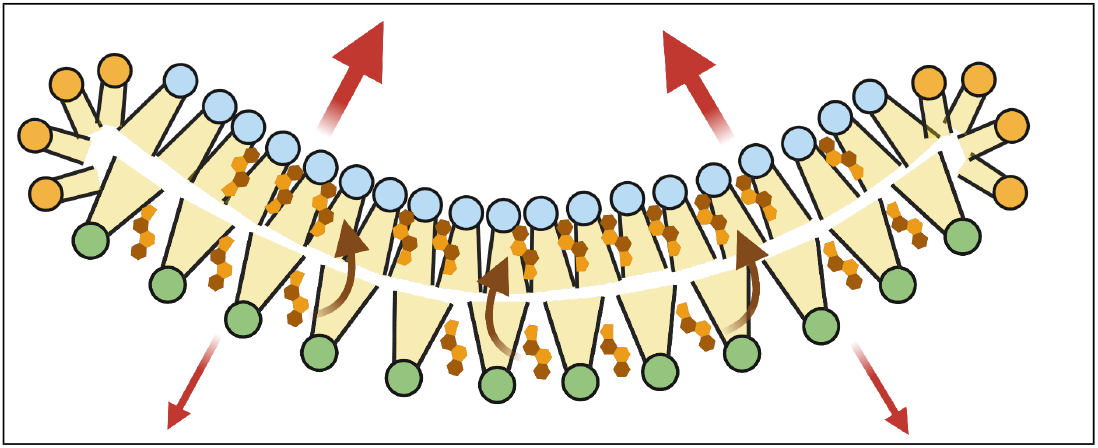

## Introduction

The composition and consecutively the organization of biological membranes gained center stage by progress in lipidomics combined with the emerging rationale that an understanding of processes at the cellular interface necessitates a molecular understanding of both protein-lipid interactions and of membrane characteristics. Apart from the functionally relevant lipid composition asymmetry across plasma membranes of eukaryotic cells^1,2^ actively maintained by e.g. lipid flippases, the cholesterol distribution between the leaflets that is determined by fast spontaneous flipping processes is still enigmatic.^3–5^

Closely connected to the cholesterol distribution between the membrane leaflets is the question of how the membrane composition affects membrane properties. Of particular interest for a number of biological questions are membrane shaping and remodeling processes, e.g. in membrane fusion,^6–8^ the role of membrane composition for the regulation of membrane protein function,^9^ the establishment of different cell identities^10^ and related in immune cell activation.^11^ In addition to specific protein-lipid interactions, driving forces include in particular collective properties such as membrane elasticity or spontaneous membrane curvature.^12^

Molecular dynamics (MD) simulations have been successfully applied in the past to shed light on typical membrane characteristics at the molecular scale.^13^ The accessible time and length scales, together with continuous improvement and extension of force fields, set MD simulations at atomistic and coarse-grained resolution into an ideal position to pinpoint the driving forces responsible for shaping (processes) at biological membranes.

The typical design of membrane simulation systems employs setups of infinite lipid bilayers with enforced periodicity across the length of the simulation system. The periodicity comes with the huge advantage that it dispenses for the need of defining membrane rims. The latter are coupled to lipid type-dependent, possibly huge rim tensions that may substantially affect membrane characteristics.^14^ In addition, periodic infinite membrane systems allow for the setup of defined asymmetries between the individual lipid leaflets that are unchanged during micro- or millisecond long simulations due to the very low lipid flipping rate.^15^ Finally, the rectangular membrane shape and the periodicity overall simplify the analysis of different observables, e.g. of the area per lipid, membrane thickness and acyl chain ordering, membrane elasticity, lipid diffusion, or of the electrostatic membrane potential.^16^ The usage of semi-isotropic pressure coupling further allows for analyzing the fluctuations of the lateral membrane area giving access to the area compressibility *K*_*A*_.

However, on the reverse side, membrane undulations are suppressed in a unit cell size-dependent way, the maximum wavelength is given by the lateral box extension, but also shorter wavelengths may be suppressed in a wavelength-dependent fashion. Additionally, the effect of membrane stress induced by unbalanced asymmetrical membrane compositions is easily blurred by the symmetrically fixed areas of both membrane leaflets: I.e., asymmetrically composed membranes setup by combining leaflets from symmetric bilayer simulations^17,18^ will typically expose an asymmetric membrane stress profile with a non-zero torque acting on the membrane.^18^ In other words, equal lipid leaflet areas do not imply a zero tension. The stress profile and connected the microscopic membrane torque are accessible from simulations of infinite membranes.^18^

However, structural effects of a net stress will not easily be uncovered in infinite membrane simulations since the overall membrane curvature vanishes across the simulation box by construction. The study of spontaneous protein-induced membrane curvature is thereby impeded as well. A setup including membrane-shaping protein(s) in both ways with respect to the membrane normal may attenuate this periodicity-dependent stress.^19^ To resolve these issues, considerable effort has been put in the development of simulation setups that allow for the study of (preset) curved membranes.^20,21^

Here, we suggest an unbiased lipid bicelle system that avoids both the periodicity-related drawbacks of infinite membrane systems and the side effects of a bicelle rim. This is achieved by defining an intermittent *overlap domain* between a *central bicelle domain* with the lipid/protein composition of interest and a surrounding *bicelle rim domain*. The overlap domain is accessible both for lipids of the central bicelle domain and of the rim domain. The latter is decorated by short-chain fatty acids. The central bicelle domain displays structural characteristics in excellent agreement to corresponding infinite bilayer systems. In addition, also the collective properties such as the membrane elasticity that is estimated from the (almost) unrestrained local curvature fluctuations are in very good agreement to infinite membrane systems. Application of the bicelle setup to a plasma membrane mimic allowed for the unbiased study of the lipid packing density within the cytoplasmic and the extracellular leaflets as well as of the debated cholesterol distribution between the layers of a plasma membrane.

## Methods

Several different membrane compositions were investigated in a lipid bicelle setup. Additionally, a set of infinite membranes was simulated to compare their characteristics with those of equally composed bicelles. For almost all setups and simulations, we employed the coarse-grained MARTINI force field.^22–24^ The different bicelle systems are listed in Table 1, additional simulations of infinite lipid bilayer systems are provided in Table S1 of the Supporting Information. An overview of some of the studied lipid types in MARTINI representation is given in Figure 1.

**Table 1:** Lipid bicelle systems. Given is the respective phospholipid to cholesterol ratio, the number of lipids and of cholesterol molecules (CHOL) within the central domain (excluding edge stabilizing rim lipids), the number of lipids within the disc interior (radius 7 nm; no rim lipids within this region), and the simulation time. Systems are labeled as bicelle systems (*d:*), without cholesterol, or high cholesterol content (*h*). Studied were pure lipid bicelles with POPC, DPPC, DOPC, and DPSM, protein-lipid bicelles with either aquaporin (AQP0) or the voltage-gated potassium channel (KvChim) embedded in POPC without cholesterol and at high cholesterol concentration, and bicelles with the asymmetric lipid composition of red blood cell plasma membranes (d:PM^0..4^).

| **System** | **Central domain:**<br>**#(lipids)/(#(CHOL))** | **Disc interior:**<br>**#(lipids)** | **Sim.time**<br>**in $\mu s$** |
| --- | --- | --- | --- |
| d:POPC | 798 | 459 | 15,15 |
| d:DPPC | 850 | 490 | 10 |
| d:DOPC | 766 | 441 | 10 |
| d:DPSM | 860 | 499 | 10 |
| d:AQP0 | 695/0 | — | 20 |
| d:AQP0<sup>h</sup> | 606/303 | — | 20 |
| d:KvChim | 688/0 | — | 20 |
| d:KvChim<sup>h</sup> | 603/302 | — | 20 |
| d:PM<sup>0</sup> | 613/438 | 388/192 | 2 |
| d:PM<sup>1</sup> | 630/451 | 397/195 | 8 |
| d:PM<sup>2</sup> | 612/439 | 388/189 | 8 |
| d:PM<sup>3</sup> | 601/430 | 383/187 | 8 |
| d:PM<sup>4</sup> | 595/426 | 380/184 | 8 |

**Figure 1:**
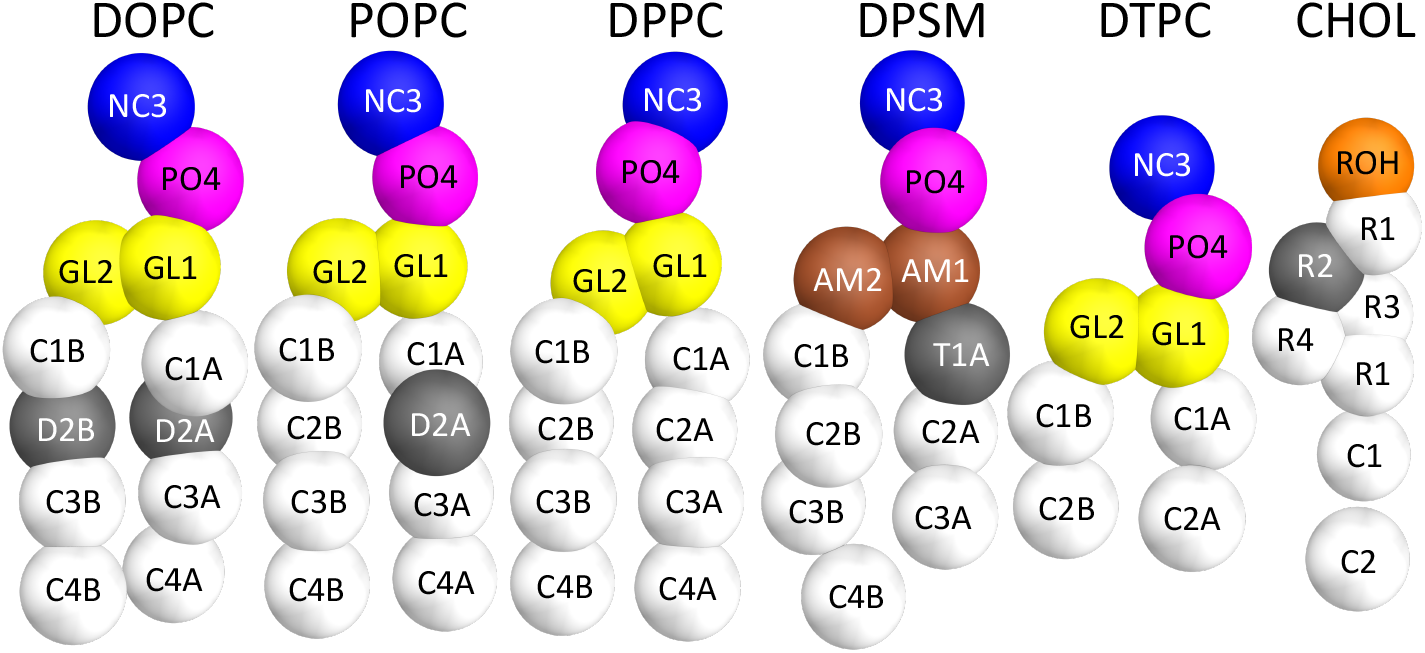
Lipids studied in monovalent lipid bicelle systems. The structure of the lipids is given in the MARTINI coarse-grained representation together with the CG atom type names. DTPC and DOPC lipids were chosen for the bicelle rim domain.

### Lipid bicelle systems

An unperturbed, monovalent phospholipid bicelle easily yields a slightly too low area per lipid and an overestimated membrane thickness that is caused by the line tension at the bicelle rims.^25^ Addition of short-chained lipids to the rims of lipid bicelles reduces the line tension and thereby overall stabilizes the system with respect to vesiculation.^14,26^ However, the study of asymmetric membranes is hampered by the largely free diffusion of lipids across the bicelle rim.

Here, we aimed for the setup of a bicelle with defined compositions of the *rim domain* and of the *central bicelle domain*. To this end, we constrained the (radial) lateral motion of lipids of the central bicelle domain to the bicelle core by a cylindrical flat bottom potential with radius R_*outer*_ = 10.5 nm (Fig. 2 a, *red line*). This domain contains the lipid-protein composition of interest. The setup also allows for an asymmetric lipid composition within this domain. In turn, lipids within the *rim domain* are constrained to the outer cylindrical region of the bicelle (inverted flat bottom potential with radius R_*inner*_ = 8.1 nm, Fig. 2 a, *blue line*). It contains lipids with both short- and long acyl chains (1,2-ditricosanoyl-snglycero-3-phosphocholine, DTPC, with short acyl chains and 1,2-dioleoyl-sn-glycero-3-phosphocholine, DOPC, with long chains). Lipids of the central domain and of the rim domain may freely exchange within the intermediate *overlap domain* for R_*inner*_ *<* **r** *<* R_*outer*_ (see Fig. 2 a,b).

**Figure 2:**
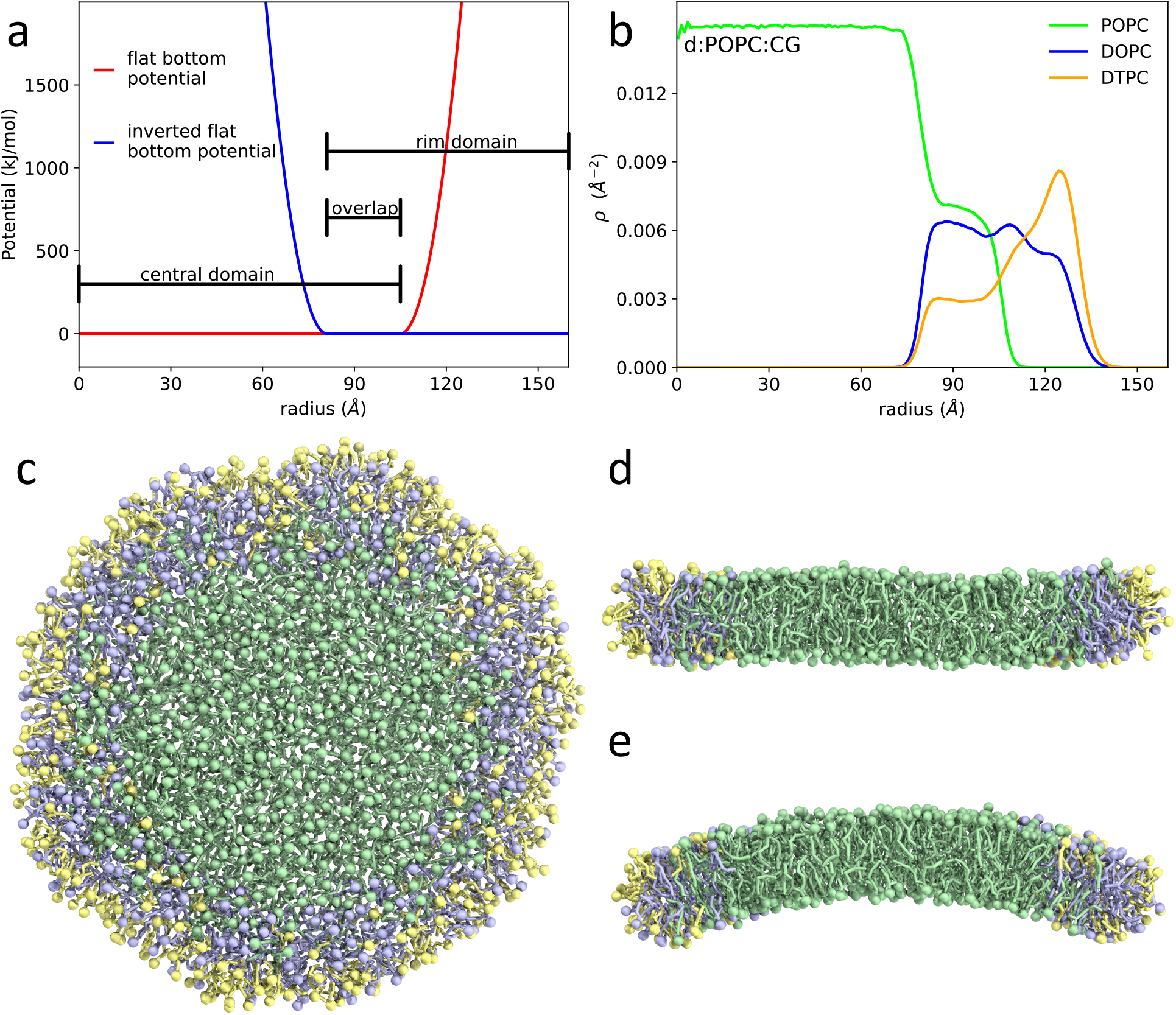
**a**. Flat bottom potential *V* (**r**) used to keep lipids away from the bicelle rim domain (*red* line) and inverted flat bottom potential (*blue*) introduced to constrain rim domain lipids to the rim and overlap zone. Averaged lipid densities per membrane area *ρ*(**r**) are shown for the d:POPC bicelle analyzed from CG simulations (**b**). **c-e**. Top view and side view cuts of lipid bicelle d:DOPC. The rim lipids are coloured in *yellow* (DTPC) and *blue* (DOPC). Lipids of the central disc domain are coloured in *green* (DOPC). Both rim and central domain lipids are found within the overlap zone.

The overlap domain forms a buffer region that allows for a smooth transition between rim and central domains and, in particular, reduces a possible bias introduced by the rim energy on the bicelle curvature, and ultimately also prevents vesiculation and diffusion of lipids of the central domain along the rim, thereby maintaining an initial lipid asymmetry in leaflet composition within the central domain. In addition, the bicelle setup may counteract moderate differences or changes in the total area of asymmetric leaflets or membranes – due to e.g. inserted asymmetric membrane proteins – by rearrangement of the rim lipids.

### Bicelle setup

The adjustment of the lipid numbers within the central bicelle domain of given size requires knowledge about the approximate area occupied by lipids within the respective membrane leaflet. To this end, simulations of symmetric infinite lipid bilayers displaying either the composition of the upper or lower leaflet were employed (for simulation details see below). For the central bicelle domain, an initial hexagonal lipid bilayer system is generated via the tool INSANE,^27^ including lipids (symmetric or asymmetric) and possibly membrane proteins (AQP and KvChim, see below). Coarse-grained protein structures for the crystal structures of AQP (PDB:2B6P^28^) and KvChim (PDB:2R9R^29^) were obtained as specified in Kluge *et al*.^25^ The target dimension of the central domain is 266 nm^2^, corresponding to a circular patch of radius ≈ 9.2 nm.

The bilayer is solvated with water and the system is minimized (500 steps steepest descent). Subsequently, the hexagonal lipid patch is placed in a larger rectangular box and re-solvated (≈ 28,000 water molecules). After 500 steps of steepest descent energy minimization, the system is equilibrated for 20 ns with a time step of 20 fs and position restraints in normal (*z*-)direction on the phosphate beads of the lipids (force constant of 1,000 kJ/mol nm^−2^) resulting in an approximately circular lipid bilayer surrounded by water.

In the next step, the bicelle rim domain is built by adding ≈ 380 long-chained DOPC lipids and ≈ 370 short-chained DTPC lipids to the rim of the central bicelle domain (surrounded by ≈ 100,000 water molecules). The DTPC and DOPC lipids spontaneously adjust to the bicelle rim during a 20 ns equilibration with position restraints in *z*-direction on the phosphate beads of the central domain lipids. Next, the position restraints were replaced with cylindrical flat bottom potentials with a force constant of 500 kJ/mol nm^−2^ on the phosphate beads of all lipids and, if present, the backbone beads of an inserted protein. The radius of the flat bottom potential was chosen to R_*outer*_ = 10.5 nm for the lipids of the central domain and to 7.1 nm for integral membrane proteins (if present). For the rim lipids DOPC and DTPC the radius of the inverted flat bottom potential was chosen to R_*inner*_ = 8.1 nm.

The simulation lengths for the pure lipid bicelle systems and the protein bicelles ranged between 10-30 *µ*s. Further details on the composition of the systems, the number of replicas, or the number of lipids within the central bicelle domain used for analysis are provided in Table 1.

Additionally, an asymmetric bicelle was set up that mimics the composition of the red blood cell plasma membrane.^1^ The lipid numbers of both leaflets were adjusted separately to match the area of the central bicelle domain, the lipid areas were taken from CG simulations of corresponding symmetric bilayers with either the composition of the extracellular plasma membrane leaflet (EL) or of the cytosolic leaflet (CL). In order to minimize the spontaneous curvature occurring in this plasma membrane bicelle (see Results Section), four additional systems were studied with lipids added or removed keeping the lipid ratios within the respective leaflets constant: d:PM^1^: 30 lipids added to the CL; d:PM^2^: 25 lipids added to CL, 25 lipids removed from EL; d:PM^3^: 10 lipids added to CL, 30 lipids removed from EL; d:PM^4^: 10 lipids added to CL, 40 lipids removed from EL. The resulting lipid compositions within a central domain of radius 7 nm averaged over the simulation time are provided in Tables 2 and 3.

**Table 2:** Plasma membrane bicelle systems. Averaged number of lipids within the extracellular leaflet of the central bicelle domain (radius 7 nm).

| System | Extracellular Leaflet |  |  |  |  |  |  |  |
| --- | --- | --- | --- | --- | --- | --- | --- | --- |
|  | PAPS | PAPE | PAPC | POPC | PIPC | DPSM | PNSM | CHOL |
| d:PM<sup>0</sup> | 1.5 | 11.7 | 18.2 | 25.1 | 53.8 | 41.0 | 64.0 | 116.3 |
| d:PM<sup>1</sup> | 1.3 | 10.7 | 17.9 | 22.9 | 51.6 | 42.4 | 60.5 | 109.5 |
| d:PM<sup>2</sup> | 1.4 | 10.2 | 17.3 | 22.6 | 48.4 | 39.9 | 59.3 | 105.0 |
| d:PM<sup>3</sup> | 1.1 | 10.7 | 16.9 | 22.7 | 48.7 | 41.2 | 60.0 | 107.6 |
| d:PM<sup>4</sup> | 1.4 | 10.2 | 16.5 | 22.3 | 47.6 | 39.7 | 58.3 | 103.9 |

**Table 3:**
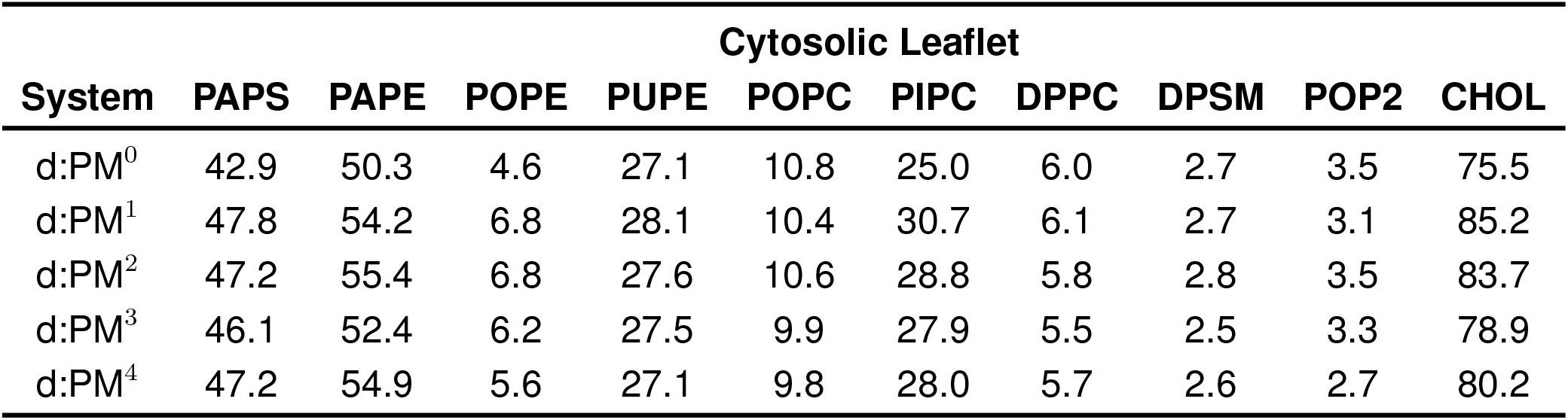
Plasma membrane bicelle systems. Averaged number of lipids within the cytosolic leaflet of the central bicelle domain (radius 7 nm).

| System | Cytosolic Leaflet |  |  |  |  |  |  |  |  |  |
| --- | --- | --- | --- | --- | --- | --- | --- | --- | --- | --- |
|  | PAPS | PAPE | POPE | PUPE | POPC | PIPC | DPPC | DPSM | POP2 | CHOL |
| d:PM<sup>0</sup> | 42.9 | 50.3 | 4.6 | 27.1 | 10.8 | 25.0 | 6.0 | 2.7 | 3.5 | 75.5 |
| d:PM<sup>1</sup> | 47.8 | 54.2 | 6.8 | 28.1 | 10.4 | 30.7 | 6.1 | 2.7 | 3.1 | 85.2 |
| d:PM<sup>2</sup> | 47.2 | 55.4 | 6.8 | 27.6 | 10.6 | 28.8 | 5.8 | 2.8 | 3.5 | 83.7 |
| d:PM<sup>3</sup> | 46.1 | 52.4 | 6.2 | 27.5 | 9.9 | 27.9 | 5.5 | 2.5 | 3.3 | 78.9 |
| d:PM<sup>4</sup> | 47.2 | 54.9 | 5.6 | 27.1 | 9.8 | 28.0 | 5.7 | 2.6 | 2.7 | 80.2 |

### Lipid bilayer systems

For comparison of membrane characteristics, simulations of (standard) infinite lipid bilayers were performed (systems denoted as e.g. *i:POPC*): For each system, two differently sized membranes were studied, a small system (labeled with index *s*) with a size comparable to the studied bicelles (800-900 lipids), and an approximately three times larger bilayer system (1,800-2,500 lipids, index *b*). Production runs had a length of 2 *µ*s with a time step of 20 fs. Details on the composition of the systems and the box size are provided in Table S1 of the Supporting Information.

### Simulation parameters

For all simulations the temperature was kept at 320 K via the v-rescale algorithm.^30^ Isotropic pressure coupling (semi-isotropic for infinite membranes) to 1 bar was employed using the Parrinello-Rahman algorithm^31,32^ with a compressibility of 3 · 10^−4^ bar^−1^. Electrostatic interactions were treated using a reaction field algorithm with a cut-off of 1.1 nm and a relative dielectric constant of 15. For the plasma membrane systems a salt concentration of 0.15 M NaCl was used, and the particle-mesh Ewald (PME) method^33^ with a relative dielectric constant of 2.5 was employed for electrostatics. A single cut-off of 1.1 nm was chosen for the Lennard-Jones interactions. Output was written every 500 ps. For all coarse-grained simulations, the MARTINI force field (version 2.2)^22–24^ was used together with the GROMACS 2020(2021) simulation package^34^.

### Analysis

The analysis of the structural and dynamical characteristics of the (protein-)lipid-bicelle systems was performed on the circular central domain of the bicelle (radius 7 nm). For infinite systems the whole system was taken into account unless otherwise specified. All errors were estimated using block averaging.

### Membrane surface

The membrane surfaces were defined based on height fields for the upper *z*^*u*^(*x, y*) and lower *z*^*l*^(*x, y*) membrane leaflets, calculated on a 2D grid in lateral direction (see also^35,36^). For each grid cell (*i, j*), the *z*_*ij*_-value was taken as the sum of the *z*-values of surrounding lipid PO4 beads (see Figure 1), weighted by a Gaussian distribution 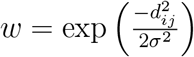. *d*_*ij*_ is the *xy*-plane distance of the PO4 bead from the respective grid point and *σ* = 0.8 nm the standard deviation. The grid spacing was chosen to ∆*x* = ∆*y* = 0.5 nm.

In presence of an integral membrane protein, the analyzed circular patches were centered around the center of mass of the protein. Furthermore cells outside the central domain of the bicelle (radius 7 nm) were neglected as well as grid cells appointed to the protein. A grid cell was appointed to the protein if a backbone bead of the protein was closer than all surrounding phosphate beads in the *xy*-plane. In order to only account for backbone beads close to the respective membrane surface, first the mean *z*-positions of the lipid PO4 beads within a 3*σ xy*-distance of the cell were calculated. Only protein backbone beads within a 0.5 nm range in *z*-direction were considered.

The bilayer surface was defined as the average of the monolayer surfaces 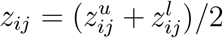 and the membrane height fields 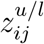 were used to calculate the local membrane normals 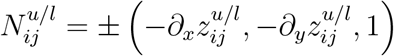.

### Membrane thickness

The height fields of the upper and lower leaflets were used to calculate the membrane thickness 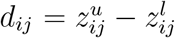. Thickness values *d* averaged over the central domain (all points for infinite systems) and all analyzed frames are provided in Table S2 of the Supporting Information.

### Local spontaneous membrane curvature

The local mean spontaneous membrane curvature was evaluated for the bilayer surface *z*_*ij*_. The mean curvature *H*_*ij*_ is given by

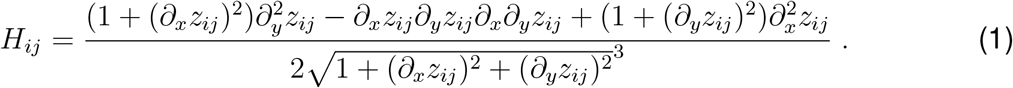

Average curvature values *H*(*t*) were computed over grid cells with a lateral distance *< R* to the bicelle center (protein center of mass for protein containing systems) at time *t*. Curvatures of infinite bilayer systems were analyzed for similar circular regions.

### Area per lipid

The area per lipid, *A*_*l*_, was calculated based on a local Voronoi tesselation for the individual lipids (see also^37^). The centers of mass of the GL1 and GL2 beads (AM1 and AM2 for DPSM) were taken as the respective lipid positions. For each lipid, Voronoi tesselation was performed considering lipids within a distance of 4 nm. First the normal is calculated for the point cloud of the positions of the selected lipids. Theses lipids were projected onto the plane perpendicular to this local membrane normal, followed by Voronoi tesselation. The area per lipid of the central lipid is taken as the area of its Voronoi cell. Values provided in Table S2 are averages over all analyzed frames and all lipids of the central domain.

### Order parameter

The order parameter *P*_2_ is defined as

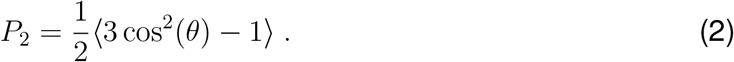

Here *θ* is the angle between the local membrane normal in the upper, respectively lower, leaflet 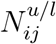 and a bond vector in the lipid tail. Each lipid was assigned the local normal of the grid cell that is closest to the center of mass of the C1A and C1B beads of the lipid (see Figure 1) and the expectation value was taken over all analyzed frames, all lipids, and all tail bonds. Order parameters for all investigated systems are provided in Table S2.

### Lipid diffusion

The Einstein relation in two dimensions is

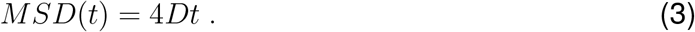

It relates the lateral mean square displacement *MSD*(*t*) after lag time *t* to the diffusion coefficient *D* (for *t* → ∞). The lateral lipid displacements were calculated for the PO4 beads within the *xy*-plane. The diffusion coefficient *D* was fitted to the mean square displacement *MSD* for lag times between 5 ns and 10 ns. For the bicelle systems, displacements were only considered for lipids within 3 nm of the bicelle center before displacement, thereby minimizing the effect of the bicelle rim. Values for the investigated systems are presented in Table S2.

### Membrane elasticity

Different approaches have been developed in the past to derive the membrane bending elasticity *κ*_*b*_ from molecular dynamics simulations. The energetics of shape deformations of membranes can be described by continuum theories like the Helfrich theory using a height field *z*(**r**)^38^ or models including more molecular details, e.g. lipid orientations as used within the lipid director field **n**(**r**).^39^ In the following, we briefly describe the orientation spectrum,^40^ undulation spectrum,^40^ and real space fluctuation methods^41,42^ employed in this work, as well as a new approximation to calculate the bending modulus directly from local fluctuations in membrane curvature.

### Orientation spectrum

The orientation spectrum method^40^ uses the lipid director field **n**(**r**) to calculate the bending modulus 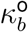. The equipartition theorem relates the power spectrum of the longitudinal component 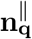 to the bending modulus 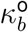:

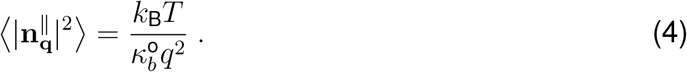

The transverse components of the lipid directors 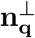 further gives access to the lipid tilt modulus *κ*_*θ*_ and the lipid twist modulus *κ*_tw_:

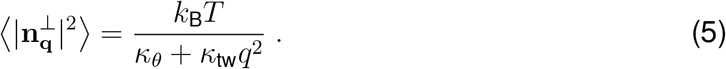

In practice, the lipid director is the vector connecting the lipid head to the lipid tail. The lipid head is the center of mass of the PO4, GL1 and GL2 (PO4, AM1, AM2 for DPSM) beads, the tail is the center of mass of the last beads of the fatty acyl chains (see Figure 1). The lipids of each leaflet were first assigned to a two dimensional grid with a grid spacing of ∼ 1 nm using the *xy*-positions of the lipid heads. The values of the lipid director fields for the upper leaflet 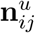 and the lower leaflet 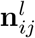 at grid position *ij* were defined as the average of the lipid directors assigned to grid point *ij* of the respective leaflet.

The discrete Fourier transforms of the monolayer director fields 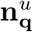 and 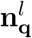 yield the bilayer director field in Fourier space 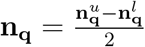 with wave vector **q**. The power spectrum of its longitudinal component 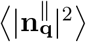 for absolute values of the wave vector *q <* 0.9 nm^−1^ was used to fit the bending modulus 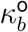 according to Eq. 4. Similarly, the tilt *κ*_*θ*_ and twist *κ*_tw_ moduli were obtained from a fit to the power spectrum of the transversal component 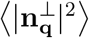 according to Eq.5 for absolute values of the wave vector *q <* 2 nm^−1^.

### Undulation spectrum

In the undulation spectrum method, the relation between the bending modulus 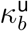 and the spectrum of the height field *z*_**q**_ is:^40^

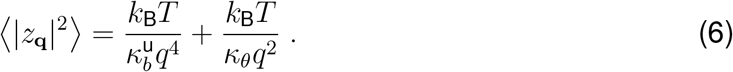

Like in the case of the lipid director fields, the lipid heads were first assigned to a twodimensional grid with a grid spacing of ∼1 nm. The height fields for the upper leaflet 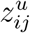 and lower leaflet 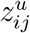 are given by the averages of the *z* positions of the assigned lipid heads. Discrete Fourier transformation yields the bilayer height field in Fourier space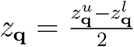 with wave vector **q**. The resulting power spectrum 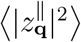 is used to obtain the bending modulus 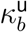 from a fit to Eq. 6 and absolute values of the wave vector *q <* 0.9 nm^−1^. The lipid tilt modulus *κ*_*θ*_ is taken from the orientation spectrum method.

### Real space fluctuations (RSF)

The geometry of the bicelle does not allow for a discrete Fourier transformation of the lipid director **n**(**r**) or height fields *z*(**r**). Therefore, additional methods that do not rely on the discrete Fourier transformation, but rather make use of local membrane properties were applied. The real space fluctuation method^41,42^ uses the distribution of splay values *S* = ∇**n** − ∇**N** between pairs of neighbouring lipids to extract the bending modulus 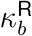. The lipid director **n** is defined as above.

The original RSF method uses an averaged surface. To include the instantaneous local surface as proposed before,^43^ here a *local* normal **N** to the grid cell that is closest to the center of mass of the C1A and C1B beads of the respective lipid was used instead. The center of mass of the C1A and C1B beads was used to calculate splay values only for neighbouring lipids (distance *<* 1.1 nm). Splay values were taken as the covariant derivative of **n** − **N** orthogonal to the local membrane normal and in the direction that is connecting the centers of mass of the C1A and C1B beads of the respective lipids. The Boltzmann distribution describes the distribution of splay values *P* (*S*):

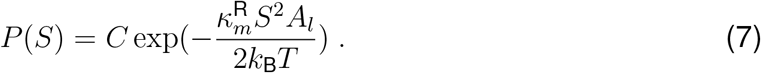

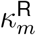 is the monolayer bending modulus and *A*_*l*_ the area per lipid. The monolayer bending modulus 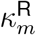 is obtained from a quadratic fit to −2*k*_B_*T* ln (*P* (*S*)) */A*_*l*_ for each monolayer and the bilayer bending modulus 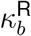 as the sum of the monolayer bending moduli.

### Local fluctuation method (LFM)

Here, we additionally derive an alternative way to calculate the bending modulus *κ*_*b*_ directly from the fluctuations in membrane curvature *H*. The membrane bending energy is given by

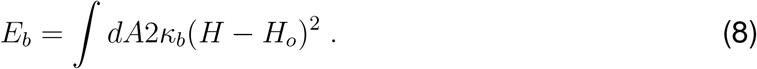

*H* is the curvature, *H*_*o*_ the spontaneous membrane curvature. In the following, the integral is approximated by a sum over the grid points *i* of a circular patch of defined size of either the lipid bicelle system or the infinite membrane systems. A symmetric bilayer with constant lipid density and thus vanishing spontaneous curvature *H*_*o*_ is assumed,

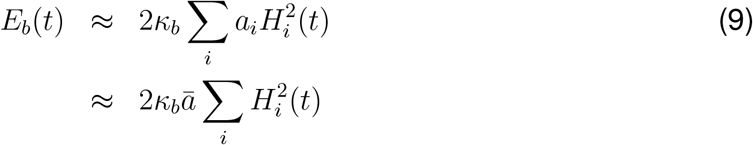

The area *a*_*i*_ of each grid point *i* is taken to be approximately constant (*a*_*i*_ = *ā* = 1*/N* · *A, N* number of grid points, *A* total area of circular patch). Next, we develop the local grid-point curvature *H*_*i*_(*t*) by introducing the average curvature of the circular patch, 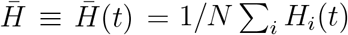:

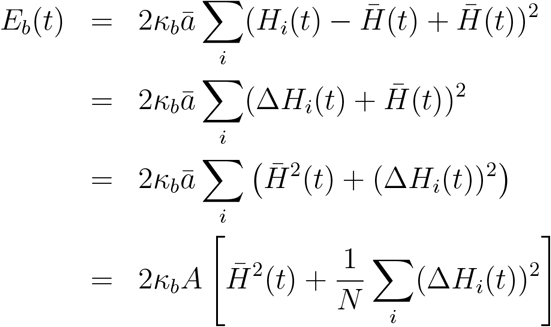

The probability for the circular patch to assume the mean curvature 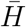 is given by:

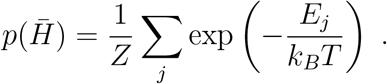

Here, *Z* is the partition function and the sum Σ_*j*_ is over all states with 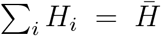 Accordingly,

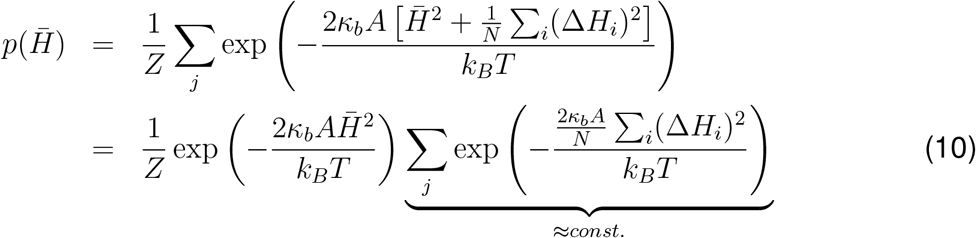

Local deviations of the curvature at each grid point ∆*H*_*i*_ from the average curvature 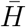 are approximately independent of the actual value of the mean curvature 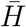, as shown in the Supporting Information Figure S1. Additionally, the degeneracy of states – that is the number of states displaying the same mean curvature (Σ_*j*_ in Eq. 10) – is independent of 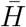. In summary, the bending elasticity may be determined from the probability density of states 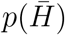. The bending elasticity is related to the variance *σ*^2^ of the approximately normally distributed 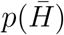by:

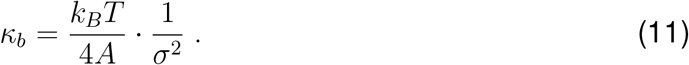

The above derivation does not explicitly consider a lipid tilt modulus. Therefore, the bending elasticities computed in this way form an upper boundary for the true elasticity.

## Results

### Lipid bicelles do not alter membrane structural characteristics

Here, we developed a novel lipid bicelle setup with a specific lipid rim domain and a central bicelle domain that are connected via an overlap domain that is accessible to lipids of both the rim and of the central domain. The latter domain is also termed *disc* in the following. The rim domain has a defined composition of DOPC and short chain lipids (DTPC) to reduce the line tension, the composition of the disc region can freely be adjusted, including asymmetric compositions. The disc region displays comparable static (area per lipid, thickness) and dynamic properties (acyl chain order) as compared to infinite membrane systems for all four analyzed lipid types differing in chain length, acyl chain saturation, and headgroup (POPC, DOPC, DPPC, DPSM; data collected in Supporting Information Table S2) thus suggesting a negligible influence of the line tension induced by the rim in the chosen setup. Finite size effects by the rim reduce the lipid diffusion coefficient within the central domain by 12-14% with respect to infinite membrane systems. Importantly, the bilayer bending modulus *κ*_*b*_ is in good agreement with experiments and simulations of infinite systems (see below).

### Short-ranged effect of channels on curvature

A *monovalent* bicellar system has been used before in the analysis of the spontaneous curvature induced by a water channel (AQP0) and a voltage-gated potassium channel (KvChim).^25^ Good agreement of the induced curvature to corresponding sorting experiments could be achieved. However, the region of the membrane less affected by the rim strain and thus suitable for the analysis of membrane properties was restricted to a circular domain of less than 5.5 nm distance from the channel center of mass (COM). Additionally, the line tension induced by the rim led to a slight compression of the membrane.

The bicelle systems developed here enables the unbiased study of the induced curvature within circular disc regions with a diameter of up to ≈ 15 nm, thereby allowing to unequivocally study the spatial range of curvature induction by transmembrane proteins. The coarse-grained simulations of the AQP0 and KvChim channels embedded in a phosphatidylcholine bicelle (POPC) show a strong positive membrane curvature (≈ +0.04 nm^−1^) induced by the potassium channel in particular between the voltage-sensing domains (circular region at a distance of 3 nm and 5 nm from protein COM) and a slight negative membrane curvature of <-0.02 nm^−1^ induced in vicinity of aquaporine (Figure 3). While the negative curvature induced by AQP0 is directly connected to its wedge-like shape, the induced positive membrane curvature close to the potassium channel is enforced by the tilted interfaces of the symmetrically arranged voltage-sensing domains at the cytosolic membrane interface of KvChim as detailed in Kluge *et al*.^25^ Interestingly, a protein-induced effect on membrane curvature is limited to a region within 1-2 nm of the protein surface.

**Figure 3:**
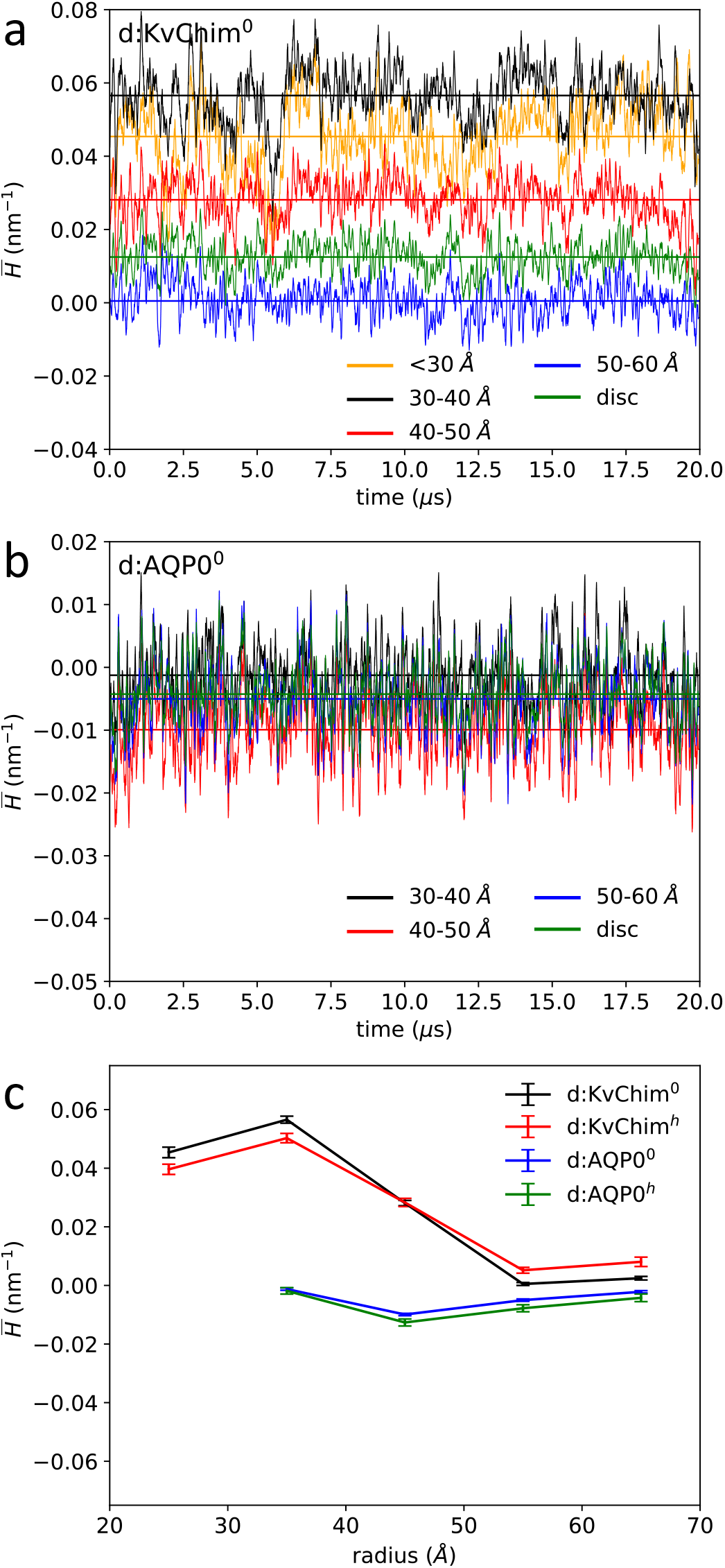
Local spontaneous membrane curvature 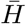 as a function of simulation time for different annuli around the center of mass of KvChim (**a**) and AQP0 (**b**), and the mean local spontaneous membrane curvature as a function of distance from the protein center of mass (**c**). Values are given as running averages with a window size of 50 ns. Straight lines mark the respective mean values.

Addition of cholesterol (33.3 mol%) did not affect the mean protein-induced membrane curvature. However, the cholesterol distribution between the layers of the membrane was strongly correlated with the disc curvature (*not shown*), with the cholesterol distribution shifted towards the inner, compressed membrane leaflet, in line with the negative spontaneous curvature observed for cholesterol both in experiments^44^ and simulations.^45,46^

### Bicelles enable unperturbed membrane fluctuations

Thermal fluctuations of membranes cause temporal local undulations. These curvature fluctuations differ substantially between the central bicelle disc domain and infinite, i.e. periodic lipid bilayers. Figure 4 compares the curvature distributions for both setups, exemplarily for a symmetric monovalent POPC system. In order to assure comparability of results, the spontaneous curvatures were averaged over circular disc regions of radius 7.0 nm for both bicelle and infinite bilayer systems. The simulation length was 10 *µ*s each. The curvature distributions could be well fitted by Gaussian distributions centered around a vanishing total curvature, differing however in the distribution widths. The full width at half maximum (FWHM) of the curvature distribution within the POPC bicelle was increased by ≈ 44% as compared to the infinite system of comparable size. Even the curvature fluctuations for a large infinite system with roughly three times more lipids was smaller by 8% as compared to the bicelle. I.e., despite similar membrane characteristics, the lipid bicelle enables significantly larger temporal fluctuations and thus curvatures that are impeded by the imposed periodic boundary conditions in infinite systems.

**Figure 4:**
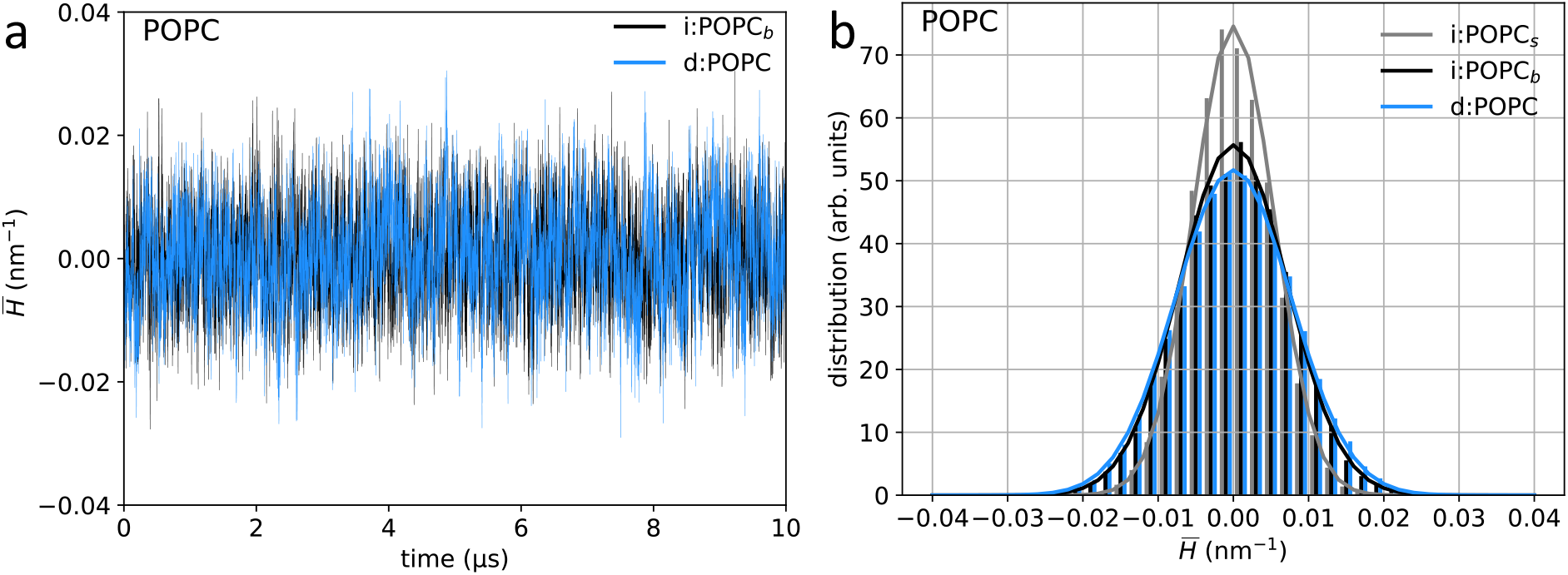
Curvature in bicelle and infinite systems: All curvature values are mean curvature values, averaged over all gridpoints within a concentric ring of radius 7.0 nm for each snapshot, both for bicelle and infinite systems. **(a)** Time development of mean curvature in a POPC bicelle (*blue*) and a large POPC bilayer extending across periodic boundaries (*black*). **(b)** Histogram of mean curvature values of a POPC bicelle (*blue*), a large (*black*) and small (*gray*) periodic POPC bilayer. The data were fitted by a Gaussian distribution (*solid lines*).

### Lipid-dependent bilayer elasticity

Most established methods to calculate the bending modulus from molecular dynamics simulations are based on the Fourier transformation of the lipid height or orientation fields.^40,47,48^ These methods can by construction not be applied to bicelle systems. Therefore, we here chose to determine the bending modulus *κ*_*b*_ from the probability density distribution 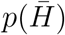 of states for the mean curvature 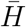 within a small circular region (’analysis spot’) of the membrane (called in the following ‘local fluctuation method’, or LFM):

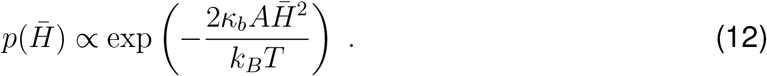

In the derivation, we assumed a vanishing spontaneous curvature *H*_*o*_ within the analysis spot, as well as a constant area *A*. A fit to the approximately normally distributed 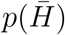 with variance *σ*^2^ allows to determine the bending elasticity:

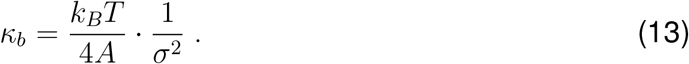

Values for the apparent *κ*_*b*_ computed in this way for different spot radii between 3.0 nm and 7.0 nm are provided in Fig. 5 for four different phospholipids. The bending moduli for small spots (radius 3.0 nm) are comparable for the different setups, i.e. the small and large infinite lipid bilayers (*gray* and *black lines* in Fig. 5) and the lipid bicelle (*blue lines*). The bending moduli for the small infinite bilayer determined via LFM (lateral box size ≈ 17 nm) quickly increase with increasing analysis spot size. I.e., membrane undulations, and thereby larger mean curvatures, are suppressed by the imposed bilayer periodicity already for circular regions with radii ≥ 4 nm. This effect is visible as well for the large infinite lipid bilayers (box size ≈ 27 nm), however shifted to larger spots with radii of at least 6-7 nm.

**Figure 5:**
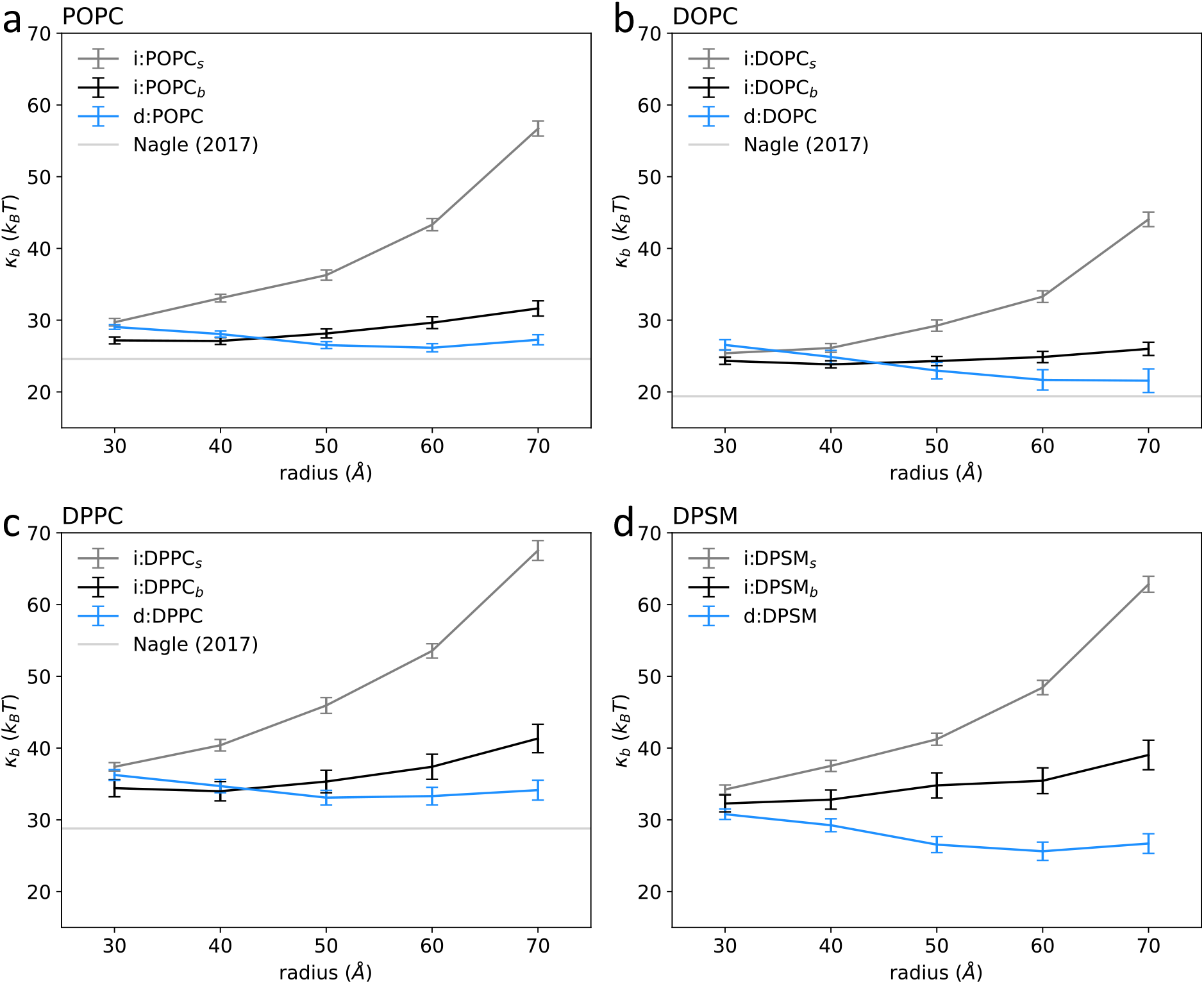
Bending elasticity *κ*_*b*_ as approximated by local fluctuation method (LFM, see Methods Section). Shown are the bending elasticities computed for differently sized circular lipid patches (i.e. for different radii) for the small (i:lipid_*s*_) and large infinite lipid bilayer systems (i:lipid_*b*_), and for the lipid bicelle systems (d:lipid) for POPC, DOPC, DPPC, and DPSM. The straight gray lines denote corresponding experimental values. ^52^

In turn, the bending modulus slightly decreases with increasing analysis spot sizes for the lipid bicelles. This decrease is attributed to the decreasing contribution of the lipid tilt to the mean curvature. Table 4 reports the bicelle bending moduli analyzed over (large) analysis spots with a radius of 7 nm, as well as the bending moduli computed using the real space fluctuation method (RSF,^42,49–51^), and determined from undulation 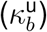 or orientation spectra 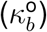. The RSF method overall yields significantly smaller values for the bending moduli compared to the other methods and will not be further considered here.

**Table 4:** Bending moduli obtained for the bicelle (d:) and for big and small infinite bilayers (i:) calculated using different methods: 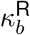, real space fluctuation method; ^42,49–51^ 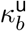, from undulation spectrum, adjusted to include lipid tilt via tilt modulus obtained from orientation fluctuations;^40,62,63^ 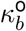, from orientation spectrum; ^40^ 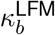, local fluctuation method, see Methods Section. The analysis for the bicelle systems was performed on the central disc domain (radius 7 nm).

| System | $\kappa_b^R$ in $k_B T$ | $\kappa_b^U$ in $k_B T$ | $\kappa_b^O$ in $k_B T$ | $\kappa_b^{LFM}$ in $k_B T$ |
| --- | --- | --- | --- | --- |
| d:POPC<sup>0</sup> | $17.1 \pm 0.1$ | — | — | $27.3 \pm 0.7$ |
| i:POPC<sub>b</sub><sup>0</sup> | $17.7 \pm 0.1$ | $24.5 \pm 0.2$ | $25.2 \pm 0.1$ | |
| i:POPC<sub>s</sub><sup>0</sup> | $17.8 \pm 0.1$ | $24.7 \pm 0.1$ | $24.5 \pm 0.1$ | |
| d:DPPC<sup>0</sup> | $23.3 \pm 0.1$ | — | — | $34.1 \pm 1.4$ |
| i:DPPC<sub>b</sub><sup>0</sup> | $24.3 \pm 0.1$ | $35.1 \pm 0.4$ | $33.7 \pm 0.3$ | |
| i:DPPC<sub>s</sub><sup>0</sup> | $24.4 \pm 0.1$ | $35.1 \pm 0.2$ | $32.7 \pm 0.1$ | |
| d:DOPC<sup>0</sup> | $14.7 \pm 0.1$ | — | — | $21.6 \pm 1.6$ |
| i:DOPC<sub>b</sub><sup>0</sup> | $15.2 \pm 0.1$ | $20.7 \pm 0.3$ | $21.9 \pm 0.2$ | |
| i:DOPC<sub>s</sub><sup>0</sup> | $15.2 \pm 0.1$ | $20.6 \pm 0.1$ | $21.1 \pm 0.1$ | |
| d:DPSM<sup>0</sup> | $23.7 \pm 0.1$ | — | — | $26.7 \pm 1.4$ |
| i:DPSM<sub>b</sub><sup>0</sup> | $24.4 \pm 0.1$ | $28.3 \pm 0.4$ | $30.1 \pm 0.3$ | |
| i:DPSM<sub>s</sub><sup>0</sup> | $24.4 \pm 0.1$ | $28.7 \pm 0.1$ | $29.7 \pm 0.1$ | |

The bending moduli determined via LFM are in very good agreement to values obtained from undulation and orientation spectra of infinite systems (exception DPSM, see Table 4). The bending modulus is largest for DPPC (32-35 *k*_*B*_*T*, depending on the particular system and analysis method) and decreases with increasing unsaturation of the lipid acyl chains (POPC 24-27 *k*_*B*_*T*, DOPC 20-22 *k*_*B*_*T*). The simulation data for *κ*_*b*_ are in overall reasonable agreement to experiment (DPPC: 28.8 *k*_*B*_*T*; ^52^ POPC: 19-25 *k*_*B*_*T*; ^52–54^ DOPC: 16-26 *k*_*B*_*T* ^52,55–61^).

### Plasma membrane bicelle

The bicelle setup introduced here is highly susceptible to changes in the composition of the central bicelle domain. In particular, a non-balanced leaflet-dependent membrane tension and resulting torque on the membrane will be reflected in a spontaneous curvature. Such a tension may easily be introduced for asymmetric membrane compositions. Here, we composed an asymmetric central bicelle domain mimicking the plasma membrane composition.^1^ Initially, the areas of the leaflets were balanced based on simulations of symmetric bilayers, reflecting either the composition of the extracellular or of the cytosolic leaflet. However, a balanced area does not imply a vanishing tension difference between the leaflets of the membrane:^18^ This initial, area-balanced setup of a plasma membrane bicelle (system d:PM^0^) led to a strong average negative spontaneous curvature within the central bicelle domain (≤ 7 nm radius) of −0.028 nm^−1^, suggesting an unbalanced membrane tension (compare Table 5).

**Table 5:** Characteristics of plasma membrane bicelles. Mean curvature 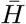, area per lipid *A*_*l*_ calculated via Voronoi tesselation for lipids and cholesterol, respectively (for extracellular^*EL*^ and cytosolic^*CL*^ leaflets), cholesterol content in extracellular and cytosolic leaflets, analyzed for central domain with radius of 7.0 nm.

| System | $\bar{H}$ in $\text{nm}^{-1}$ | $A_l^{EL}$ in $\text{nm}^2$ | $A_l^{CL}$ in $\text{nm}^2$ | Chol<sup>EL</sup> | Chol<sup>CL</sup> |
| --- | --- | --- | --- | --- | --- |
| d:PM<sup>0</sup> | -0.0278 | 0.533/0.448 | 0.682/0.542 | 35.1 mol% | 30.4 mol% |
| d:PM<sup>1</sup> | -0.0003 | 0.558/0.466 | 0.654/0.521 | 34.6 mol% | 31.0 mol% |
| d:PM<sup>2</sup> | 0.0054 | 0.565/0.472 | 0.650/0.517 | 34.5 mol% | 30.7 mol% |
| d:PM<sup>3</sup> | -0.0046 | 0.555/0.464 | 0.663/0.526 | 34.8 mol% | 30.3 mol% |
| d:PM<sup>4</sup> | 0.0034 | 0.563/0.471 | 0.657/0.521 | 34.7 mol% | 30.4 mol% |

Counterintuitively, *addition* of lipids to the positively curved cytosolic domain (system d:PM^1^), possibly combined with removal of lipids from the negatively curved extracellular leaflet (d:PM^2^-d:PM^4^) was required to balance the tension and thus the spontaneous curvature. Addition and removal of lipids was in all cases chosen such that the relative composition of the leaflets was unchanged. The significantly reduced or vanishing curvature observed upon addition of lipids to the positively curved cytosolic leaflet is likely coupled to the negative spontaneous curvature not only of phosphatidylethanolamines (PE headgroups) and cholesterol, but also of phosphatidylcholine (PC) lipids with unsaturated acyl chains.^64^ I.e., the positive curvature of the cytosolic leaflet of the starting plasma membrane bicelle is reduced by addition of lipids with negative spontaneous curvature. The decrease of curvature is further connected to a decrease of the area per lipid within the cytosolic leaflet, and an area increase for the lipids of the extracellular leaflet (see Table 5).

The equilibrated plasma membrane bicelle with vanishing spontaneous curvature (d:PM^1^), i.e. a balanced stress profile, is characterized by a cholesterol density within the extracellular leaflet that is larger by 28% compared to the cytosolic leaflet (0.648 nm^−2^ vs. 0.505 nm^−2^, ≈ 35 mol% cholesterol vs. ≈ 30 mol% cholesterol). Interestingly, the plasma membrane bicelle is significantly softer compared to a pure POPC bilayer, with an upper bound for the bending modulus of ≈ 21 *k*_*B*_*T* (*κ*_*b*_ = 29 *k*_*B*_*T* for POPC, using the local fluctuation method), despite the comparably high lipid density within the leaflets.

## Discussion

A setup for a lipid bicelle system is presented that displays similar structural and dynamical characteristics to large infinite bilayer setups. The introduction of flat bottom potentials in opposite directions for lipids within the central bicelle domain and for lipids forming the rim, and usage of lipids with short acyl-chains within the rim minimize the membrane edge tension and additionally prevent the exchange of lipids between the central domain leaflets via the bicelle edge. Since the bicelle system bypasses the boundary conditions of infinite membrane MD simulations it is ideally suited for the unbiased study of membrane curvature fluctuations and of protein- or lipid-induced spontaneous membrane curvatures. The comparative analysis of differently sized periodic infinite bilayer systems with that of lipid bicelles revealed a strong influence of periodic system size on the curvature distribution already on length scales below half the box dimension. In addition, we show that the membrane elasticity can reliably be estimated from the analysis of bicelle curvature distributions.

Results obtained for a plasma membrane-like asymmetric lipid composition suggest that the lipid-shape driven spontaneous curvature of membranes dominates over area-driven spontaneous curvature.^12^ In our example, the curvature of a negatively curved plasma membrane may be balanced by *removing* lipids from the negatively curved and densely packed extracellular leaflet and/or by *addition* of lipids to the positively curved cytosolic membrane leaflet (see sketch in Fig. 6). The opposite would be expected for membrane curvature driven by area. Our results for the direction of the spontaneous curvature, and its change upon addition/removal of lipids suggest that both plasma membrane leaflets are characterized by an inherent negative curvature. Addition of lipids, while keeping the ratio of lipids constant, increases the negative curvature of the respective leaflet, however to a different degree for the two leaflets of the plasma membrane.

**Figure 6:**
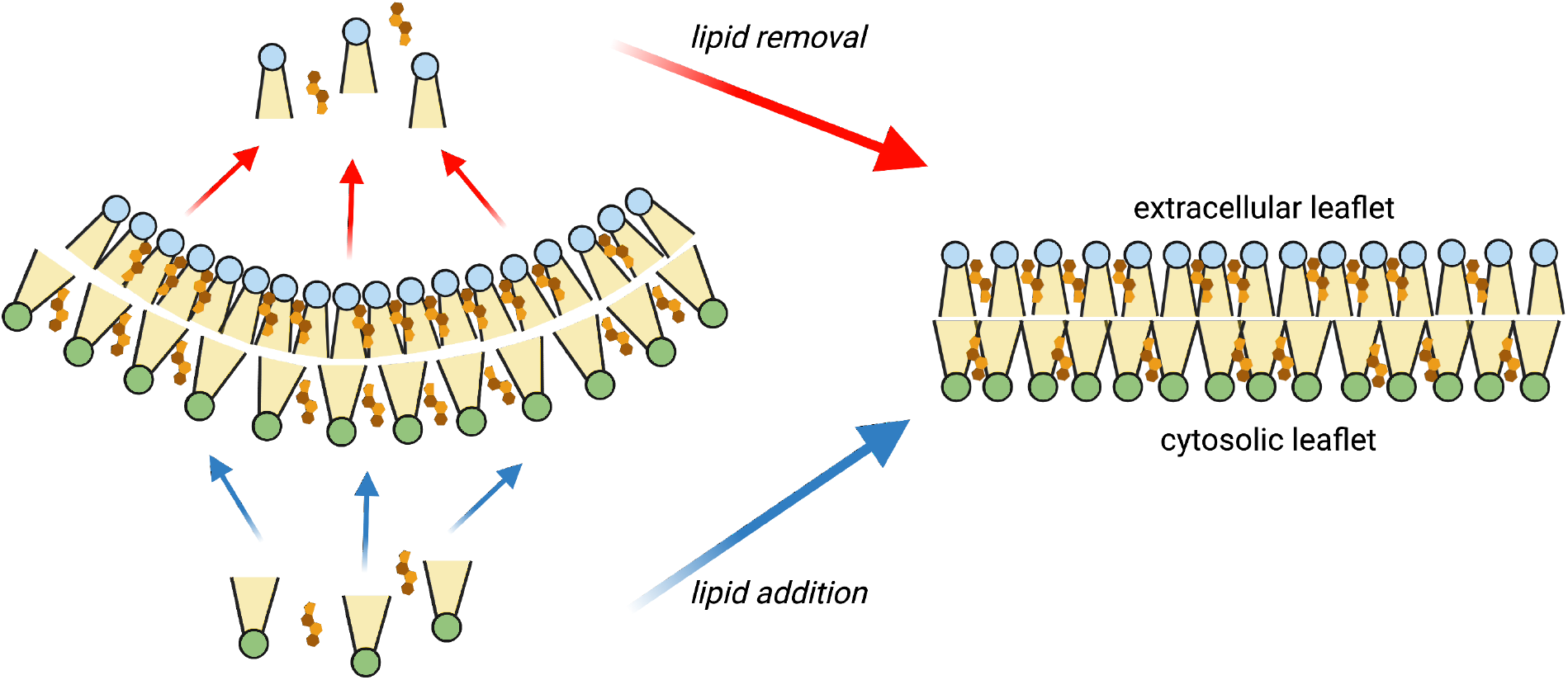
Sketched is a negatively curved plasma membrane that may be straightened by both addition of lipids to the cytosolic leaflet and/or removal of lipids from the extracellular leaflet. The composition of the individual leaflets is kept unchanged.

The flipping of cholesterol molecules may result in tensionless leaflets as shown before for infinite bilayers;^18^ Still, a substantial torque may arise that results in spontaneous curvature^18^ as directly observed here for the plasma membrane bicelle system. This spontaneous curvature is accompanied by significant changes of the phospholipid and cholesterol areas that are directly accessible in the bicelle system (compare Table 5). For a balanced plasma membrane composition (system d:PM^1^) – i.e. a torque-free membrane with vanishing spontaneous curvature – the cholesterol density is increased by 28% within the extracellular leaflet that displays an overall rather dense packing with an average phospholipid area of only ≈ 0.56 nm^2^.

## Supporting Information Available

Additional information on simulation systems, membrane characteristics, and local curvature fluctuations (PDF)

## Acknowledgement

The authors gratefully acknowledge the compute resources and support provided by the Erlangen Regional Computing Center (RRZE) and the Erlangen National High-Performance Computing Center (NHR@FAU). RAB acknowledges support by the German Science Foundation (DFG) within the SFB1027, *Physical Modeling of Non-Equilibrium Processes in Biological Systems* (project C6).

## References

(1) Lorent, J. H.; Levental, K. R.; Ganesan, L.; Rivera-Longsworth, G.; Sezgin, E.; Doktorova, M.; Lyman, E.; Levental, I. Plasma membranes are asymmetric in lipid unsaturation, packing and protein shape. Nat. Chem. Biol. 2020, 16, 644–652.

(2) Doktorova, M.; Symons, J. L.; Levental, I. Structural and functional consequences of reversible lipid asymmetry in living membranes. Nat. Chem. Biol. 2020, 16, 1321–1330.

(3) Liu, S.-L.; Sheng, R.; Jung, J. H.; Wang, L.; Stec, E.; O’Connor, M. J.; Song, S.; Bikkavilli, R. K.; Winn, R. A.; Lee, D.; Baek, K.; Ueda, K.; Levitan, I.; Kim, K.-P.; Cho, W. Orthogonal lipid sensors identify transbilayer asymmetry of plasma membrane cholesterol. Nat. Chem. Biol. 2017, 13, 268–274.

(4) Courtney, K. C.; Fung, K. Y.; Maxfield, F. R.; Fairn, G. D.; Zha, X. Comment on ‘Orthogonal lipid sensors identify transbilayer asymmetry of plasma membrane cholesterol’. Elife 2018, 7, e38493.

(5) Courtney, K. C.; Pezeshkian, W.; Raghupathy, R.; Zhang, C.; Darbyson, A.; Ipsen, J. H.; Ford, D. A.; Khandelia, H.; Presley, J. F.; Zha, X. C24 sphingolipids govern the transbilayer asymmetry of cholesterol and lateral organization of model and live-cell plasma membranes. Cell Rep. 2018, 24, 1037–1049.

(6) McMahon, H. T.; Kozlov, M. M.; Martens, S. Membrane curvature in synaptic vesicle fusion and beyond. Cell 2010, 140, 601–605.

(7) Risselada, H. J.; Grubmüller, H. How SNARE molecules mediate membrane fusion: recent insights from molecular simulations. Curr. Opin. Struct. Biol. 2012, 22, 187–196.

(8) Lipowsky, R. Remodeling of Membrane Shape and Topology by Curvature Elasticity and Membrane Tension. Adv. Biology 2022, 6, 2101020.

(9) Levental, I.; Lyman, E. Regulation of membrane protein structure and function by their lipid nano-environment. Nat. Rev. Mol. Cell Biol. 2022, doi:10.1038/s41580–022–00524–4.

(10) D’Angelo, G.; La Manno, G. The lipotype hypothesis. Nat. Rev. Mol. Cell Biol. 2022, doi:10.1038/s41580–022–00556–w.

(11) Lühr, J. J. et al. Maturation of monocyte-derived DCs leads to increased cellular stiffness, higher membrane fluidity, and changed lipid composition. Front. Immunol. 2020, 11, 590121.

(12) Hossein, A.; Deserno, M. Spontaneous curvature, differential stress, and bending modulus of Asymmetric Lipid Membranes. Biophys. J. 2020, 118, 624–642.

(13) Marrink, S. J.; Corradi, V.; Souza, P. C. T.; Ingólfsson, H. I.; Tieleman, D. P.; Sansom, M. S. P. Computational modeling of realistic cell membranes. Chem. Rev. 2019, 119, 6184–6226.

(14) de Joannis, J.; Jiang, F. Y.; Kindt, J. T. Coarse-grained model simulations of mixed-lipid systems: composition and line tension of a stabilized bilayer edge. Langmuir 2006, 22, 998–1005.

(15) Gonzalez, M. A.; Bresme, F. Membrane–ion interactions modify the lipid flip-flop dynamics of biological membranes: A molecular dynamics study. J. Phys. Chem. B 2020, 124, 5156–5162.

(16) Pluhackova, K.; Kirsch, S. A.; Han, J.; Sun, L.; Jiang, Z.; Unruh, T.; Böckmann, R. A. A critical comparison of biomembrane force fields: Structure and dynamics of model DMPC, POPC, and POPE bilayers. J. Phys. Chem. B 2016, 120, 3888–3903.

(17) Doktorova, M.; Weinstein, H. Accurate in silico modeling of asymmetric bilayers based on biophysical principles. Biophys. J. 2018, 115, 1638–1643.

(18) Miettinen, M. S.; Lipowsky, R. Bilayer membranes with frequent flip-flops have tensionless leaflets. Nano Lett. 2019, 19, 5011–5016.

(19) Rabe, M.; Aisenbrey, C.; Pluhackova, K.; de Wert, V.; Boyle, A. L.; Bruggeman, D. F.; Kirsch, S. A.; Böckmann, R. A.; Kros, A.; Raap, J.; Bechinger, B. A Coiled-Coil peptide shaping lipid bilayers upon fusion. Biophys. J. 2016, 111, 2162–2175.

(20) Yesylevskyy, S.; Khandelia, H. EnCurv: Simple technique of maintaining global membrane curvature in molecular dynamics simulations. J. Chem. Theory Comput. 2021, 17, 1181–1193.

(21) Larsen, A. H. Molecular dynamics simulations of curved lipid membranes. Int. J. Molec. Sci. 2022, 23, 8098.

(22) Marrink, S. J.; Risselada, H. J.; Yefimov, S.; Tieleman, D. P.; De Vries, A. H. The MARTINI force field: Coarse grained model for biomolecular simulations. J. Phys. Chem. B 2007, 111, 7812–7824.

(23) Monticelli, L.; Kandasamy, S. K.; Periole, X.; Larson, R. G.; Tieleman, D. P.; Marrink, S.-J. The MARTINI coarse-grained force field: extension to proteins. J. Chem. Theory Comput. 2008, 4, 819–834.

(24) de Jong, D. H.; Singh, G.; Bennett, W. D.; Arnarez, C.; Wassenaar, T. A.; Schafer, L. V.; Periole, X.; Tieleman, D. P.; Marrink, S. J. Improved parameters for the MARTINI coarse-grained protein force field. J. Chem. Theory Comput. 2013, 9, 687–697.

(25) Kluge, C.; Pöhnl, M.; Böckmann, R. A. Spontaneous local membrane curvature induced by transmembrane proteins. Biophys. J. 2022, 121, 671–683.

(26) Pannuzzo, M.; Raudino, A.; Böckmann, R. A. Peptide-induced membrane curvature in edge-stabilized open bilayers: A theoretical and molecular dynamics study. J. Chem. Phys. 2014, 141.

(27) Wassenaar, T. A.; Ingólfsson, H. I.; Böckmann, R. A.; Tieleman, D. P.; Marrink, S. J. Computational lipidomics with insane: A versatile tool for generating custom membranes for molecular simulations. J. Chem. Theory Comput. 2015, 11, 2144–2155.

(28) gonen, t.; Cheng, Y.; Sliz, P.; Hiroaki, Y.; Fujiyoshi, Y.; Harrison, S. C.; Walz, T. Lipid– protein interactions in double-layered two-dimensional AQP0 crystals. Nature 2005, 438, 633–638.

(29) Long, S. B.; Tao, X.; Campbell, E. B.; MacKinnon, R. Atomic structure of a voltage-dependent K+ channel in a lipid membrane-like environment. Nature 2007, 450, 376–382.

(30) Bussi, G.; Donadio, D.; Parrinello, M. Canonical sampling through velocity rescaling. J. Chem. Phys. 2007, 126, 014101.

(31) Parrinello, M.; Rahman, A. Polymorphic transitions in single crystals: A new molecular dynamics method. J. Appl. Phys. 1981, 52, 7182–7190.

(32) Nosé, S.; Klein, M. Constant pressure molecular dynamics for molecular systems. Mol. Phys. 1983, 50, 1055–1076.

(33) Darden, T.; York, D.; Pedersen, L. Particle mesh Ewald: An Nlog(N) method for Ewald sums in large systems. J. Chem. Phys. 1993, 98, 10089–10092.

(34) Abraham, M. J.; Murtola, T.; Schulz, R.; Páll, S.; Smith, J. C.; Hess, B.; Lindahl, E. GROMACS: High performance molecular simulations through multi-level parallelism from laptops to supercomputers. SoftwareX 2015, 1, 19–25.

(35) Allen, W. J.; Lemkul, J. A.; Bevan, D. R. GridMAT-MD: a grid-based membrane analysis tool for use with molecular dynamics. J. Comput. Chem. 2009, 30, 1952–1958.

(36) Gapsys, V.; De Groot, B. L.; Briones, R. Computational analysis of local membrane properties. J. Comput.-Aided Mol. Des. 2013, 27, 845–858.

(37) Buchoux, S. FATSLiM: a fast and robust software to analyze MD simulations of membranes. Bioinformatics 2017, 33, 133–134.

(38) Helfrich, W. Elastic properties of lipid bilayers: Theory and possible experiments. Z. Naturforsch., C, J. Biosci. 1973, 28, 693–703.

(39) Watson, M. C.; Penev, E. S.; Welch, P. M.; Brown, F. L. Thermal fluctuations in shape, thickness, and molecular orientation in lipid bilayers. J. Chem. Phys. 2011, 135.

(40) Watson, M. C.; Brandt, E. G.; Welch, P. M.; Brown, F. L. Determining biomembrane bending rigidities from simulations of modest size. Phys. Rev. Lett. 2012, 109, 1–5.

(41) Khelashvili, G.; Kollmitzer, B.; Heftberger, P.; Pabst, G.; Harries, D. Calculating the bending modulus for multicomponent lipid membranes in different thermodynamic phases. J. Chem. Theory Comput. 2013, 9, 3866–3871.

(42) Johner, N.; Harries, D.; Khelashvili, G. Curvature and lipid packing modulate the elastic properties of lipid assemblies: Comparing HII and lamellar phases. J. Phys. Chem. Lett. 2014, 5, 4201–4206.

(43) Allolio, C.; Haluts, A.; Harries, D. A local instantaneous surface method for extracting membrane elastic moduli from simulation: Comparison with other strategies. Chem. Phys. 2018, 514, 31–43.

(44) Chen, Z.; Rand, R. P. The influence of cholesterol on phospholipid membrane curvature and bending elasticity. Biophys. J. 1997, 73, 267–276.

(45) Baoukina, S.; Ingólfsson, H. I.; Marrink, S. J.; Tieleman, D. P. Curvature-induced sorting of lipids in plasma membrane tethers. Adv. Theory Simul. 2018, 1, 1800034.

(46) Kollmitzer, B.; Heftberger, P.; Rappolt, M.; Pabst, G. Monolayer spontaneous curvature of raft-forming membrane lipids. Soft Matter 2013, 9, 10877–10884.

(47) Goetz, R.; Gompper, G.; Lipowsky, R. Mobility and elasticity of self-assembled membranes. Phys. Rev. Lett. 1999, 82, 221.

(48) Lindahl, E.; Edholm, O. Mesoscopic undulations and thickness fluctuations in lipid bilayers from molecular dynamics simulations. Biophys. J. 2000, 79, 426–433.

(49) Khelashvili, G.; Harries, D. How cholesterol tilt modulates the mechanical properties of saturated and unsaturated lipid membranes. J. Phys. Chem. B 2013, 117, 2411–2421.

(50) Khelashvili, G.; Johner, N.; Zhao, G.; Harries, D.; Scott, H. L. Molecular origins of bending rigidity in lipids with isolated and conjugated double bonds: The effect of cholesterol. Chem. Phys. Lipids 2014, 178, 18–26.

(51) Doktorova, M.; Harries, D.; Khelashvili, G. Determination of bending rigidity and tilt modulus of lipid membranes from real-space fluctuation analysis of molecular dynamics simulations. Phys. Chem. Chem. Phys. 2017, 19, 16806–16818.

(52) Nagle, J. F. Experimentally determined tilt and bending moduli of single-component lipid bilayers. Chem. Phys. Lipids 2017, 205, 18–24.

(53) Arriaga, L. R.; López-Montero, I.; Monroy, F.; Orts-Gil, G.; Farago, B.; Hellweg, T. Stiffening effect of cholesterol on disordered lipid phases: A combined neutron spin echo + dynamic light scattering analysis of the bending elasticity of large unilamellar vesicles. Biophys. J. 2009, 96, 3629–3637.

(54) Mell, M.; Moleiro, L. H.; Hertle, Y.; Fouquet, P.; Schweins, R.; López-Montero, I.; Hellweg, T.; Monroy, F. Bending stiffness of biological membranes: What can be measured by neutron spin echo? Eur. Phys. J. E 2013, 36, 1–13.

(55) Pan, J.; Mills, T. T.; Tristram-Nagle, S.; Nagle, J. F. Cholesterol perturbs lipid bilayers nonuniversally. Phys. Rev. Lett. 2008, 100, 1–4.

(56) Chakraborty, S.; Doktorova, M.; Molugu, T. R.; Heberle, F. A.; Scott, H. L.; Dzikovski, B.; Nagao, M.; Stingaciu, L. R.; Standaert, R. F.; Barrera, F. N.; Katsaras, J.; Khelashvili, G.; Brown, M. F.; Ashkar, R. How cholesterol stiffens unsaturated lipid membranes. Proc. Natl. Acad. Sci. USA 2020, 117, 21896–21905.

(57) Kucerka, N.; Tristram-Nagle, S.; Nagle, J. F. Structure of fully hydrated fluid phase lipid bilayers with monounsaturated chains. J. Membr. Biol. 2005, 208, 193–202.

(58) Jablin, M. S. Tilt-dependent analysis of diffuse X-ray scattering from oriented stacks of fluid phase lipid bilayers. Ph.D. thesis, Carnegie Mellon University, 2015.

(59) Gracià, R. S.; Bezlyepkina, N.; Knorr, R. L.; Lipowsky, R.; Dimova, R. Effect of cholesterol on the rigidity of saturated and unsaturated membranes: Fluctuation and electrodeformation analysis of giant vesicles. Soft Matter 2010, 6, 1472–1482.

(60) Sorre, B.; Callan-Jones, A.; Manneville, J.-B.; Nassoy, P.; Joanny, J.-F.; Prost, J.; Goud, B.; Bassereau, P. Curvature-driven lipid sorting needs proximity to a demixing point and is aided by proteins. Proc. Natl. Acad. Sci. USA 2009, 106, 5622–5626.

(61) Rawicz, W.; Olbrich, K. C.; McIntosh, T.; Needham, D.; Evans, E. Effect of chain length and unsaturation on elasticity of lipid bilayers. Biophys. J. 2000, 79, 328–339.

(62) Levine, Z. A.; Venable, R. M.; Watson, M. C.; Lerner, M. G.; Shea, J. E.; Pastor, R. W.; Brown, F. L. Determination of biomembrane bending moduli in fully atomistic simulations. J. Am. Chem. Soc. 2014, 136, 13582–13585.

(63) Chaurasia, A. K.; Rukangu, A. M.; Philen, M. K.; Seidel, G. D.; Freeman, E. C. Evaluation of bending modulus of lipid bilayers using undulation and orientation analysis. Phys. Rev. E 2018, 97, 1–12.

(64) Venable, R. M.; Brown, F. L. H.; Pastor, R. W. Mechanical properties of lipid bilayers from molecular dynamics simulation. Chem. Phys. Lipids 2015, 192, 60–74.

