## Supporting Information for "Lipid bicelles in the study of biomembrane characteristics"

**Table S1:** Lipid bilayer simulation systems. Given are the total number of lipids, the number of lipids within the central domain (excluding edge stabilizing rim lipids), the number of lipids within the central bicelle domain (radius 7 nm; no rim lipids within this region), and the simulation time. Studied were (infinite) monovalent lipid bilayers with POPC, DPPC, DOPC, and DPSM, as well as symmetric infinite bilayers displaying either the composition of the extracellular leaflet of plasma membrane (i:PM$^{EL}$) or that of the cytosolic leaflet (i:PM$^{CL}$).

| System | Bilayer size: #Lipids | Analysis domain size: #Lipids | Sim.time in $\mu s$ | Box length in $nm$ |
|---|---|---|---|---|
| i:POPC$_b$ | 2312 | 463 | 10 | 27.7 |
| i:POPC$_s$ | 882 | 462 | 10 | 17.1 |
| i:DPPC$_b$ | 2450 | 494 | 4 | 27.6 |
| i:DPPC$_s$ | 968 | 494 | 10 | 17.4 |
| i:DOPC$_b$ | 2312 | 444 | 10 | 28.3 |
| i:DOPC$_s$ | 882 | 443 | 10 | 17.5 |
| i:DPSM$_b$ | 2450 | 500 | 4 | 27.5 |
| i:DPSM$_s$ | 968 | 500 | 10 | 17.3 |
| i:PM$^{EL}$ | 262/178 | - | 1 | 10.3 |
| i:PM$^{CL}$ | 234/174 | - | 1 | 10.4 |

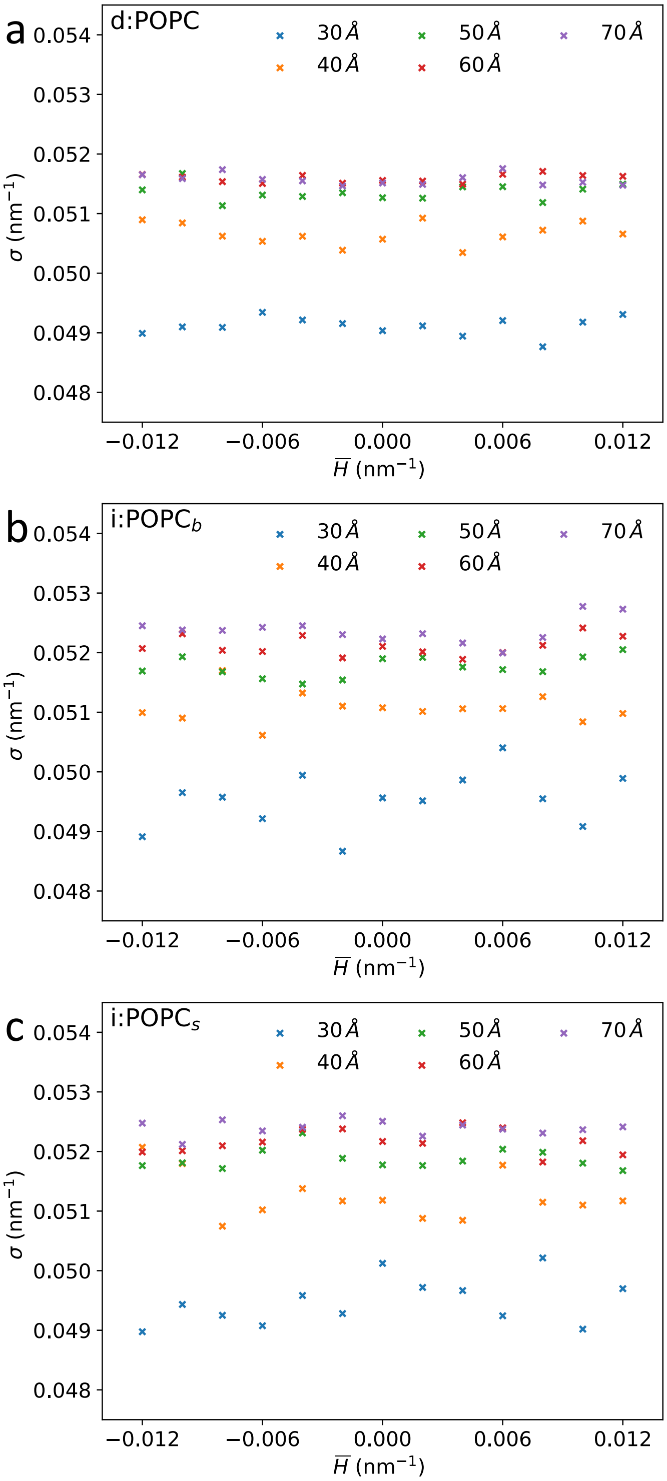


**Figure S1:** Local curvature deviations $\Delta H_i(t) = H_i(t) - \bar{H}(t)$ from the mean curvature $\bar{H}(t)$ are independent of $\bar{H}(t)$: Shown are the standard deviations $\sigma$ of the distributions of local deviations $\Delta H_i$ as a function of the mean curvature $\bar{H}$ for different circular membrane domains for the POPC bicelle (**a**), large (**b**) and small (**c**) infinite POPC bilayers.

**Table S2:** Membrane thickness $d$, area per lipid $A_l$, averaged tail order parameter $P_2$ and diffusion coefficient $D$. Properties of the lipid bicelle systems were analyzed within the central bicelle domain (radius 7 nm), those of the infinite systems for all lipids.

| **System** | $d$ in nm | $A_l$ in nm$^2$ | $P_2$ | $D$ in $1 \times 10^{-6}$ cm$^2$ s$^{-1}$ |
|---|---|---|---|---|
| d:POPC | $3.88 \pm 0.009$ | $0.681 \pm 0.00025$ | $0.528 \pm 0.0002$ | $0.793 \pm 0.006$ |
| i:POPC$_b$ | $3.88 \pm 0.002$ | $0.672 \pm 0.00006$ | $0.535 \pm 0.00004$ | $0.927 \pm 0.004$ |
| i:POPC$_s$ | $3.88 \pm 0.002$ | $0.671 \pm 0.00004$ | $0.536 \pm 0.00006$ | $0.917 \pm 0.004$ |
| d:DPPC | $4.08 \pm 0.015$ | $0.637 \pm 0.0005$ | $0.607 \pm 0.0004$ | $0.724 \pm 0.007$ |
| i:DPPC$_b$ | $4.07 \pm 0.003$ | $0.628 \pm 0.00004$ | $0.614 \pm 0.00007$ | $0.854 \pm 0.005$ |
| i:DPPC$_s$ | $4.07 \pm 0.003$ | $0.627 \pm 0.00005$ | $0.614 \pm 0.00009$ | $0.837 \pm 0.005$ |
| d:DOPC | $3.76 \pm 0.009$ | $0.710 \pm 0.0004$ | $0.473 \pm 0.0006$ | $0.863 \pm 0.008$ |
| i:DOPC$_b$ | $3.77 \pm 0.008$ | $0.701 \pm 0.00002$ | $0.480 \pm 0.00004$ | $1.00 \pm 0.005$ |
| i:DOPC$_s$ | $3.76 \pm 0.002$ | $0.700 \pm 0.00003$ | $0.481 \pm 0.00005$ | $0.975 \pm 0.006$ |
| d:DPSM | $3.78 \pm 0.023$ | $0.629 \pm 0.0009$ | $0.648 \pm 0.0004$ | $0.807 \pm 0.006$ |
| i:DPSM$_b$ | $3.76 \pm 0.003$ | $0.621 \pm 0.00004$ | $0.652 \pm 0.00005$ | $0.928 \pm 0.007$ |
| i:DPSM$_s$ | $3.75 \pm 0.002$ | $0.621 \pm 0.00004$ | $0.652 \pm 0.00005$ | $0.923 \pm 0.005$ |
